# 3D Interferometric Lattice Light-Sheet Imaging

**DOI:** 10.1101/2020.08.27.266999

**Authors:** Bin Cao, Guanshi Wang, Jieru Li, Alexandros Pertsinidis

## Abstract

Understanding cellular structure and function requires live-cell imaging with high spatio-temporal resolution and high detection sensitivity. Direct visualization of molecular processes using single-molecule/super-resolution techniques has thus been transformative. However, extracting the highest-resolution 4D information possible from weak and dynamic fluorescence signals in live cells remains challenging. For example, some of the highest spatial resolution methods, e.g. interferometric (4Pi) approaches^1-6^ can be slow, require high peak excitation intensities that accelerate photobleaching or suffer from increased out-of-focus background. Selective-plane illumination (SPIM)^7-12^ reduces background, but most implementations typically feature modest spatial, especially axial, resolution. Here we develop 3D interferometric lattice light-sheet (3D-iLLS) imaging, a technique that overcomes many of these limitations. 3D-iLLS provides, by virtue of SPIM, low light levels and photobleaching, while providing increased background suppression and significantly improved volumetric imaging/sectioning capabilities through 4Pi interferometry. We demonstrate 3D-iLLS with axial resolution and single-particle localization precision down to <100nm (FWHM) and <10nm (1σ) respectively. 3D-iLLS paves the way for a fuller elucidation of sub-cellular phenomena by enhanced 4D resolution and SNR live imaging.

---

Interferometric (4Pi) approaches^1-6^ achieve the highest 3D spatial resolution and single-particle localization precision to date, <100nm and <10nm along *z* respectively. Point-scanning 4Pi imaging^1^, with focused excitation and confocal detection, features the most efficient 3D PSF in terms of reduced side-lobes as well as background reduction. However, point-scanning is limited in temporal resolution when imaging large fields, and short pixel dwell times require high peak excitation intensities that often accelerate photobleaching. Wide-field interferometric setups could in principle achieve faster imaging at reduced peak intensities. However such setups typically have been implemented in an epi-illumination configuration^2-6^, which suffers from increased out-of-focus background. Applications have thus been limited to relatively sparse and bright cellular structures with low background. There are many systems where structures of interest might comprise only ∼10 molecules^3,13^. Such structures are difficult to visualize in the presence of often overwhelming cellular background^13^. An approach that can simultaneously achieve high 3D resolution and localization precision and reduced background/photo-bleaching would greatly advance our capabilities to visualize molecular structures and motions at nanometer scales, in the high-background, crowded intracellular milieu.

Selective plane-illumination (light-sheet) approaches illuminate only a thin slice through the sample, overcoming many of the limitations of epi-illumination. In the simplest implementations, a single plane is illuminated with an excitation beam that is perpendicular to the detection optics. A conventional choice for creating the selective plane illumination profile is a Gaussian beam, which can be projected by a separate excitation objective lens mounted perpendicular to the detection lens^10^, or reflected by a microfabricated cantilever mirror mounted on an excitation objective lens opposed to the detection lens^9^. In either case, a trade-off between the thickness of the light sheet and the effective field of view due to diffraction needs to be considered and a sweet spot is chosen depending on the requirements of the sample studied. To overcome the constraints due to diffraction of Gaussian beams, non-diffracting beams, such as Bessel^8,11^ or Airy^12^ beams can be used. Both a single beam^11^ and an array of Bessel beams^8^ can be scanned to create a light sheet that is thinner than what achieved with a Gaussian beam. However, the non-negligible excitation side-lobes away from the main illumination plane introduce excess background and unnecessary photo-bleaching at out-of-focus parts of the sample that are not imaged. Selective plane illumination based on bound 2D optical lattices, lattice light-sheet (LLS) illumination, can suppress the side lobes while maintaining the non-diffracting property and the thin profile of the light sheet^7^. LLS microscopy demonstrated combined low photo-toxicity, low photo-bleaching and low background, well-suited to live cell imaging studies. In conventional LLS imaging with a single 0.7 NA excitation lens and a single 1.1 NA detection lens, a 240nm×240nm×380nm resolution has been achieved^7^, while for single-particle localization applications, typical localization precisions^7,14^ are ∼20nm in *xy* and ∼45nm in *z*. This *z* performance is significantly lower that what can be achieved by interferometric methods, but unfortunately, possibly due to the constrains of the dual opposed objective lens geometry, it has been challenging to implement selective-plane illumination approaches in interferometric setups, to further reduce background and increase the achievable 3D resolution.

Here we report an interferometric imaging method for highly-sensitive live-cell imaging that replaces the original epi-illumination scheme with a selective-plane illumination scheme based on optical lattices (LLS illumination). This novel 3D interferometric lattice light-sheet (3D-iLLS) imaging method, achieves a more confined detection volume than conventional LLS microscopy with a single detection objective lens, and thus less out-of-focus background and higher signal-to-noise ratio. The reduced background, higher photon utilization efficiency and the higher optical sectioning capabilities of 3D-iLLS enable visualizing weak sub-cellular structures. We demonstrate an achievable *z* resolution <100nm (FWHM) and a localization precision <10nm (1σ), both a factor of ∼4× improvement compared to conventional LLS.

To better understand and optimize 3D-iLLS microscopy, we develop a numerical simulation pipeline (Supplementary Note; Supplementary Figure 1) for calculating the resulting 3D PSFs based on electromagnetic vector-field calculations. The overall 3D-iLLS PSFs show distinct profiles compared to conventional LLS microscopy (Supplementary Figure 2). In 3D-iLLS with constructive emission interference, the PSF exhibits a maximum centered at the common focus of the two objectives, with two additional visible side lobes along the *z*-axis. When emission interferes destructively, the intensity maxima are symmetrically positioned along the *z*-axis away from the focal plane, with two less pronounced side lobes. In both cases, the volume occupied by the overall PSFs for 4Pi detection is ≈2× less than for 2Pi detection, indicating reduced background and thus higher sensitivity when imaging single molecules and other faint objects at the focal plane.

Our 3D-iLLS microscope (Figure 1**a**; Supplementary Figure 3) uses two 1.1 NA opposed detection lenses, in a previously described^3^ interferometric cavity arrangement. For imaging, the interferometer is tuned to the zero-path-length position, resulting in constructive/destructive interference at the two ports of the beam-splitter. An third 0.7 NA lens, orthogonal to the two opposed detection lenses, delivers the LLS excitation. We calibrate the experimental 3D-iLLS PSF using 40-nm fluorescent beads (Figure 1**b**). The experiment calibrations recapitulate our numerical calculations, featuring the expected modulated PSF with a ∼100-140nm FWHM central lobe, thus demonstrating successful implementation of the desired 3D-iLLS optical properties. Further optimization should be possible (Supplementary Figure 4), by correcting residual aberrations in both the detection and excitation paths that most likely limit the performance of the current instrument.

**Figure 1.**
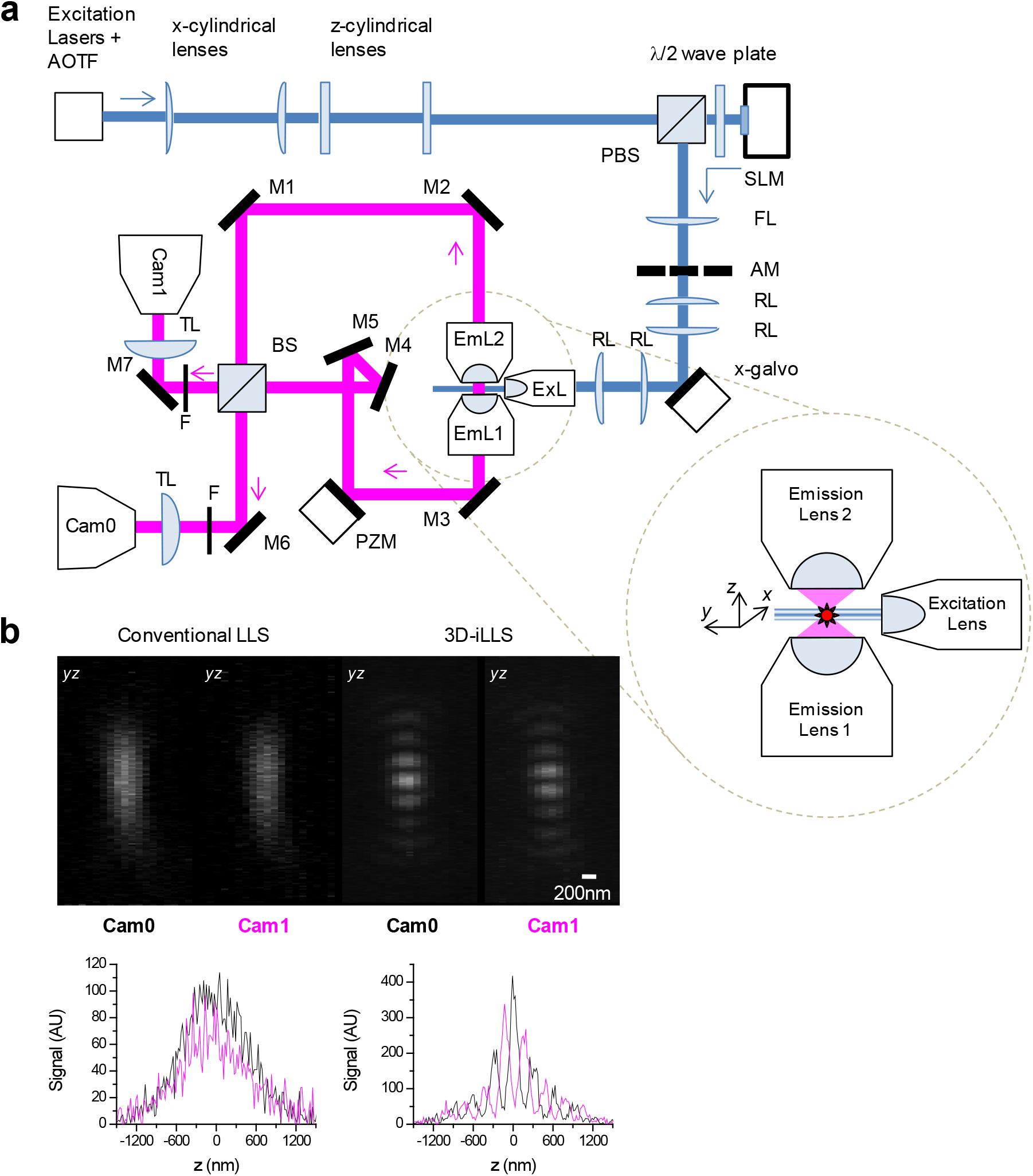
Principles of 3D-iLLS, optical setup schematic and comparison of PSF properties of 3D-iLLS vs. conventional LLS. **a**, Schematic of experimental setup. ExL: excitation les; EmL1,2: emission detection lenses. M1-7: mirrors. BS: non-polarizing beam-splitter. PZM: piezo-electric mount (phase shifter). Cam0,1: sCMOS detection cameras. TL: tube lens. F: emission filters. AOTF: Acousto-optic tunable filter. PBS: polarizing beam-splitter. SLM: spatial light modulator. AM: annular mask. FL: Fourier-transform lens. RL: relay lenses. Inset: geometry and coordinate system between lenses. **b**, Experimental calibration of the conventional LLS and 3D-iLLS PSFs, using 40nm fluorescent beads. Excitation: 642nm; emission filter: 700/70m.

To test the performance of 3D-iLLS with cellular samples we visualized single mRNAs in human osteosarcoma cells (U-2 OS)^13^. Single mRNAs are tagged with PP7 phage derived stem-loops (24×PP7) and visualized with a tandem-dimer phage coat protein fused to Halo-tag (tdPCP-Halo) and staining with a JF-646 Halo-tag ligand. 3D-iLLS adequately resolved single mRNAs, even in tight clusters where conventional LLS failed (Figure 2**a**). We further quantified the axial resolution by measuring the *z* profile of single mRNAs. Whereas the obtained FWHM resolution is 444±80 nm (mean±S.D., *n*=4) for conventional LLS, the 3D-iLLS z profiles yield a FWHM of 96±10 nm (mean±S.D., *n*=4) (Figure 2**b**). These results demonstrate a ≈4× axial resolution improvement of 3D-iLLS compared to conventional LLS.

**Figure 2.**
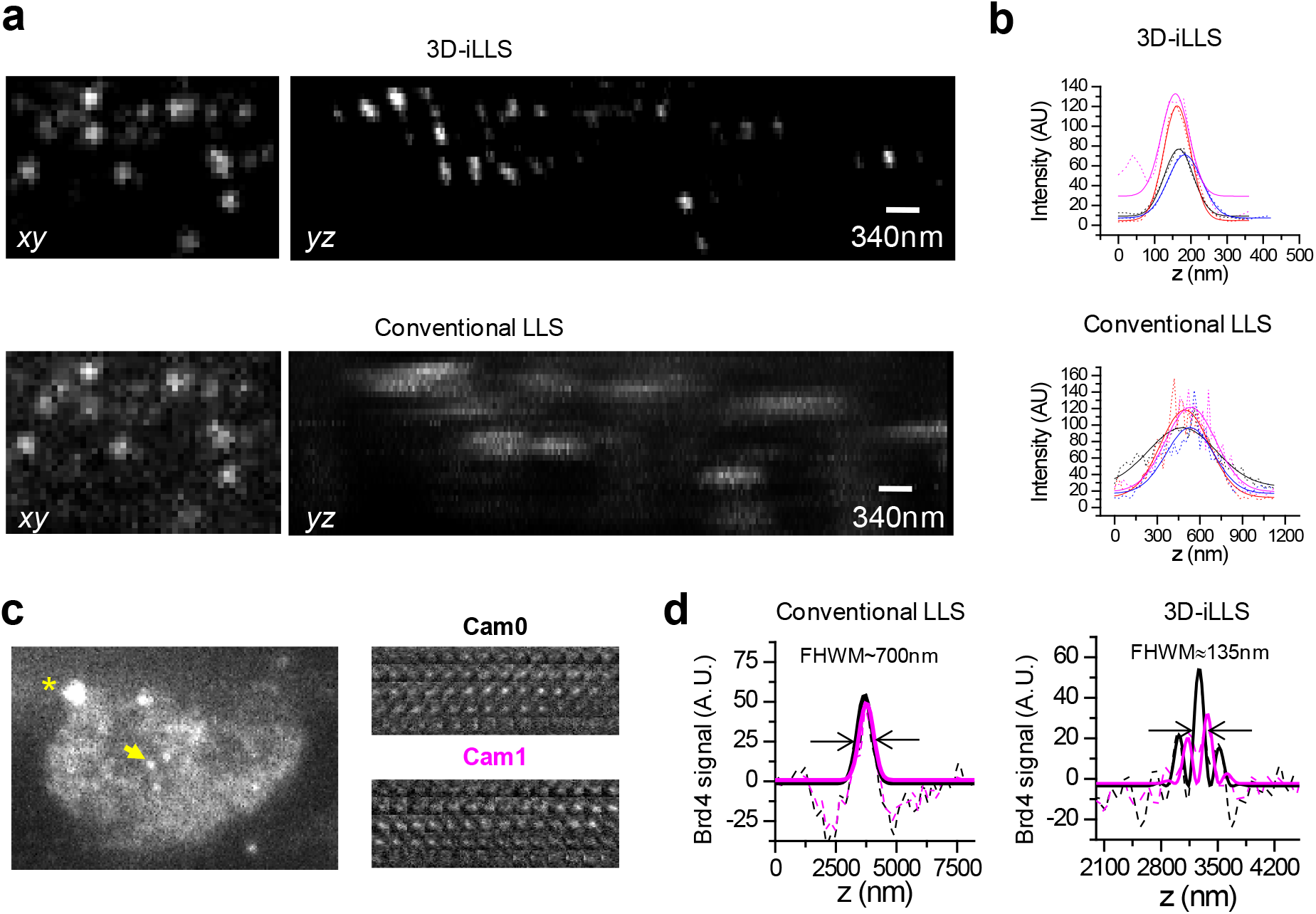
3D-iLLS outperforms conventional LLS in 3D sub-cellular imaging. **a**, Comparison of 3D-iLLS vs. conventional LLS for imaging 24×PP7 mRNAs tagged with tdPCP-Halo-JF646 in fixed U-2 OS cells. *xy* and *yz* maximum projections projections. **b**, *z* profiles of single mRNAs (dashed lines) and 1D Gaussian peak fits (solid lines), showing increased axial resolution for 3D-iLLS vs. conventional LLS. **c**, 3D-iLLS image of Brd4 clusters in live mESCs. A single *z* slice from the raw data is shown. Asterisk marks a bead fiducial. Insets show individual *z* slices for the cluster indicated by the yellow arrow (3D-iLLS data, 20nm *z* steps). **d**, *z* profiles (background-subtracted) of single Brd4 clusters imaged with conventional LLS and 3D-iLLS. Solid lines: non-linear least-squares fit to 1D Gaussian peaks for conventional LLS and to equations of the form 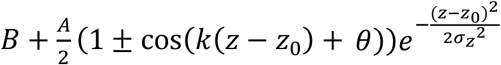 for 3D-iLLS, for Cam0 and Cam1 respectively. Data in **c, d** are raw data without deconvolution. Data in **a, b** are shown after deconvolution.

To demonstrate the performance of our 3D-iLLS setup in live-cell imaging, we imaged mouse embryonic stem cells (mESCs) that are engineered with a SNAP-tag knocked into the endogenous *Brd4* locus^13^. We had previously shown that Brd4 forms foci containing ∼15 tagged molecules at the enhancers of the *Pou5f1* and *Nanog* genes^13^. Our 3D-iLLS imaging shows multiple Brd4 clusters throughout the nucleus of mESCs (Figure 2**c**), suggesting extensive Brd4 clustering at mESC enhancers. 3D-iLLS resolves Brd4 clusters with increased resolution compared to conventional LLS (Figure 2**d**). 3D-iLLS can also localize the center-of-mass of Brd4 clusters in the reconstructed cellular volumes with ≈10nm *z* localization precision (Supplementary Figure 5). These results highlight the capabilities of 3D-iLLS for live cell imaging.

Although objects like Brd4 clusters can be localized in 3D-iLLS reconstructed cellular volumes, imaging sequential optical sections by *z* scanning compromises speed. Certain applications such as single-molecule localization-based imaging and single-particle tracking require fast 3D coordinate determination, in a narrow range within the focal plane, with sub-diffraction precision. Such localization measurements can greatly benefit from the reduced background in selective-plane illumination schemes, but conventional LLS microscopy with a single detection objective and astigmatism-based axial detection could only achieve ∼40-50nm *z* localization precision^7, 14, 15^. We reason that the ∼10× more efficient photon utilization efficiency^3^ of interferometric vs. astigmatism-based localization and the 2× higher SNR of 3D-iLLS vs. conventional LLS should push the 3D nanometer localization precision to the sub-10 nm regime.

To harness 3D-iLLS for improved *z* localization precision, we implement modulation interferometry^3^, a method that previously achieved ∼1-2nm *z* localization precision. We extract the *z* position by dynamically modulating the length of one of the interferometer arms and measuring the phase of the ensuing intensity modulation (Figure 3**a** and Supplementary Figure 6**a**). Previously modulation inteferometry relied on the coherence of two counter-propagating excitation beams. Here, we instead rely of the coherence of the emitted fluorescent photons. Importantly, for emission interference, the two ports of the interferometer beam splitter are shifted by ≈90° relative to each other, thus enabling measuring two phases simultaneously, on the two detection cameras respectively. We take advantage of this effect to improve the temporal resolution of modulation interferometry by 2-fold (Figure 3**b** and Supplementary Figure 6**b**). Using the combined phases from Cameras 0 and 1, with a 6-phase or 4-phase modulation cycle, our 3D-iLLS setup achieves ≈2nm and ≈8nm *z* localization precision respectively (Figure 3**c,d** and Supplementary Figure 6**c**,**d**). This “open loop” performance, without any active stabilization, indicates short-term mechanical stability of the 3D-iLLS design in the <10nm regime. 3D-iLLS and modulation interferometry also enabled successfully tracking the 3D movement of single 24×PP7 mRNAs tagged with tdPCP-Halo-JF646 in the cytoplasm of live U-2 OS cells (Figure 3**e**). These results illustrate how 3D-iLLS can also be exploited for dynamic 3D single-particle tracking in live cells.

**Figure 3.**
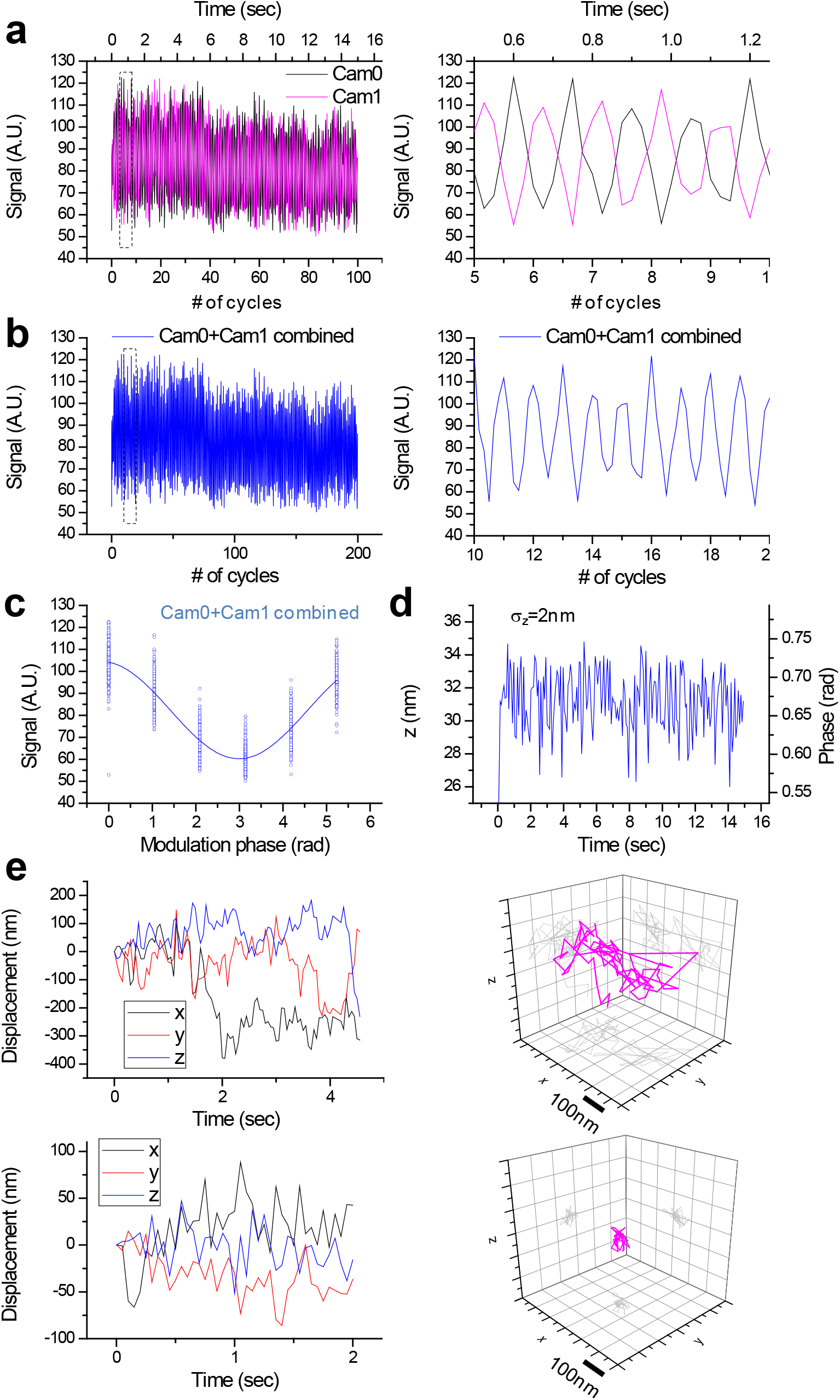
3D-iLLS and modulation interferometry enable improved axial localization and 3D single-particle tracking. **a-d**, Axial localization performance with 3D-iLLS and 6-phase modulation interferometry. **a**, Signals from a 40nm bead on Cameras 0 and 1, over 100 6-step modulation cycles. The piezoelectric phase shifter is stepped in 121.67nm increments, corresponding to 0°, 60°, 120°, 180°, 240°and 300° relative phases. Right panel: zoom-in of the dotted region in the left panel, illustrating the anti-correlated signal modulation of Cam0 vs. Cam1. **b**, Signals from Cam0 and Cam1 are combined into a single modulation cycle, doubling the temporal resolution. Right panel: zoom-in of the dotted region in the left panel. **c**, Superposition of all 200 modulation cycles by collapsing the *x* axis in the interval [0-2π), showing excellent stability and reproducibility of the setup. Solid line: fit to a sine wave. **d**, Extracted phase and *z* coordinate, showing σ_*z*_≈2nm r.m.s. localization precision. **e**, 3D tracking of single mRNAs in live U-2 OS cells using 3D-iLLS and 4-phase modulation interferometry. The piezoelectric phase shifter is stepped in 182.5nm increments, corresponding to 0°, 90°, 180° and 270° relative phases. Cam0 and cam1 signals are combined for extracting *z* coordinates. See also Supplementary Figure 6. Top panels: an mRNA molecule undergoing mostly random Brownian motion; bottom panels: an mRNA molecule confined to a volume of ∼20 nm r.m.s. radius in 3D.

Our results establish 3D-iLLS as a versatile technique with improved volumetric imaging of crowded cellular samples. Further developments are possible. 3D-iLLS and modulation interferometry^3^ could also enhance 3D localization-based super-resolution imaging, with the reduced background being particularly useful when imaging densely labeled samples^14^. Adaptive optics^16^ to correct for system and sample induced aberrations could further optimize the interferometric PSF properties, reducing the axial extend of the detection and excitation PSFs and well as better matching the detection PSFs to increase coherence. These optimizations could further reduce the axial side-lobes of the interferometric PSF, with significant gains for deconvolution applications^17^. Moreover, fast adaptive optics, such as deformable mirrors, in the detection path would also enable real-time corrections of chromatic aberrations, an important prerequisite for multi-color 3D-iLLS imaging. Finally, our design can be further augmented by introducing additional excitation lenses^18^ and implementing structured illumination microscopy (SIM) schemes^4, 19^, for eventual live-cell 3D-iLLS-SIM at sub-100nm isotropic 3D resolution.

## Methods

### 3D-iLLS microscope setup

The 3D-iLLS setup is built on an actively stabilized vibration isolation platform (TMC, Stacis iX) inside a temperature-controlled room. The 3D-iLLS microscope consists of a LLS excitation path orthogonal to the optical axis of two opposed detection lenses in a 4Pi interferometric arrangement. We use a custom 26.7× 0.7 NA excitation objective lens (Special Optics, 54-10-7) and two 25× 1.1 NA detection objective lenses (Nikon, MRD77225).

#### Detection path

The first detection lens is mounted on a 3D flexure stage driven by differential micrometers (Thorlabs, MBT616D). The second detection lens is mounted on a 3D flexure stage configured with differential micrometers and additional closed-loop piezoelectric actuators (Thorlabs, MAX301). An additional 50mm-travel stage driven by a stepper motor (Thorlabs, LNR50S) is used for coarse positioning of the second detection lens. The interferometric detection path is similar to our previous modulation interferometry setup^3^, including a motorized platform for coarse path-length scanning and a piezoelectric phase shifter for fine-tuning/fast modulation (Physik Instrumente, S-303.CD with E-709.CHG controller). The fluorescence beams from the two detection lenses interfere at a non-polarizing beam splitter, and the light from the two exit ports is filtered using quad notch filters (Semrock, StopLine NFO3-405/488/561/635E-25) and emission filters (Chroma, ET700/75m; selectable using a filter wheel; Thorlabs, FW103H), before it is imaged onto two sCMOS cameras (Hamamatsu, C11440-22CU) with two f=50cm achromatic lenses. Final magnification is ≈100nm/pixel.

#### Excitation path

The excitation path is built based on the originally reported LLS design^7^. An ATOF device (AA OPTO-ELECTRONIC, ATFnC-400.650-TN and MPDS8C driver) facilitates on-off switching of a CW laser beam (642nm; MPB Communications, 2RU-VFL-P-500-642-B1R or 2RU-VFL-P-2000-642-B1R) and modulating laser power. An elliptical lens telescope reshapes the excitation beams before they are spatially modulated by a combination of a phase-only spatial-light-modulator (SLM; Forth Dimension Displays, QXGA-3DM), an achromatic quarter-wave plate and a polarizing beam splitter cube. The modulated beam is Fourier-Transformed with a lens and the unmodulated light is blocked using a custom annular mask (Photo Sciences, custom design). The resulting 2D optical lattice is dithered using an *x*-axis galvanometer (Thorlabs, GVS001), placed a plane conjugate to the Back-Focal-Plane (BFP) of the excitation lens. Sample- and BFP-conjugate cameras (Thorlabs, DCC1545M and Edmund EO-0312M) are used for inspection of the 2D lattice pattern.

#### Instrument control and synchronization

Instrument control and synchronization are achieved with a custom Lab-VIEW (National Instruments, 2015 64-bit) application and an FPGA-based real-time hardware system (National Instruments, PCIe-7852R LX50).

#### Sample mounting and sample cell

To position electron microscopy grids in the space between the 3 objective lenses, we machine a pincher-grip sample holder out of a stainless-steel rod. The sample holder is mounted on a rotation mount and pitch-adjustable kinematic adapter (Thorlabs, RSP05 and TPA01), which are further mounted on a 3D nanopositioning stage (Physik Instrumente, P-733.3DD with E-727.3CDA controller) and an *xyz* micrometer-driven translation stage (Thorlabs, LNR25M). The sample is immersed from the top into a cubic chamber machined out of stainless-steel (Supplementary Figure 3).

Three of the horizontal faces of the chamber contain openings for the detection and excitation lenses. Silicone sheets (∼100μm thick) stretch over each lens and provide sealing. The fourth horizontal face of the sample holder contains a glass window for observing the interior of the chamber.

#### Live-cell imaging

For live-cell imaging, the temperature of the three objectives and the sample cell is regulated to ∼37°C, using resistive heaters and thermistor sensors. A separate temperature controller (Thorlabs, TC200) regulates the temperature of each objective lens, as well as a heating plate at the base and a cover plate at the top of the sample cell respectively. The exact temperature set-points are determined empirically, while the temperature of the media inside the sample cell is verified with an independent out-of-loop probe. The top cover plate is further used for delivering a mixture of N_2_, CO_2_ and O_2_ gases, with flow-rates controlled by three independent mass-flow controllers (Omega Engineering).

### Cell sample preparation

#### EM grid preparation, bead deposition and cell seeding

Cells were cultured on gold electron microscopy (EM) grids with Formvar films (Ted-Pella, 01703G). For mESC cells, EM grids were pre-coated with 5mg/ml laminin (BioLamina LN511) at 37°C overnight in a humidified 5% CO2 incubator. For U-2 OS cells, EM grids were pre-coated with collagen (Sigma C8919) at 37°C overnight in a humidified 5% CO2 incubator. In both cases, the EM grids were coated with the Formvar film facing up. After coating, EM grids were briefly rinsed with 1×PBS solution and flipped over for adding fluorescent bead fiducials. 0.1 μm TetraSpeck beads (ThermoFisher Scientific T7279) were diluted 1:300 with 1×PBS and MgCl_2_ was added to the diluted beads to a final concentration of ∼ 0.1M. Diluted beads were then deposited onto EM grids for 10min at 37°C and removed, rinsed briefly with 1×PBS. EM grids were then flipped over with the Formvar film facing up and placed in glass bottom microwell dishes (MatTek, P35G-1.5-14-C), ready for cell seeding. Cells were trypsinized, counted and ∼ 0.3 million cells were seeded in the appropriate media.

#### Cell culture and staining

All cell cultures were maintained at 37°C, in 5% v/v CO2 atmosphere, in a humidified incubator. Brd4 OMG cells were cultured with +2i media with 400 µg/ml G418 (Sigma G8168-10ML) on 0.1% gelatin-coated dish, at 37°C in a humidified 5% CO2 incubator. +2i media contain D-MEM (Thermo Fisher Scientific 10313021), 15% fetal bovine serum (Gemini Bio 100-500), 0.1 mM 2-mercaptoethanol (Thermo Fisher Scientific 21985023), 2 mM L-alanyl-L-glutamine (Thermo Fisher Scientific 35050079), 1× MEM nonessential amino acids (Thermo Fisher Scientific 11140076), 1000 U/mL LIF (Millipore ESG1107), 3uM CHIR99021 (Millipore 361559) and 1 uM PD0325901 (Axon Medchem 1408). Before imaging, cells were seeded with -2i media plus 400 µg/ml G418. For SiR staining, cells were labeled with 0.3μM SiR-BG for 10 min, at 37°C, followed by three times rinsing with new media.

CMV clone 5 cells were maintained in McCoy’s 5A media without phenol-red (GE Healthcare SH30200.01), supplemented with 10% fetal bovine serum (Gemini Bio, 100-500), 1× Non-essential Amino Acids Solution (Thermo Fisher Scientific 11140050), 1mM Sodium Pyruvate (Thermo Fisher Scientific 11360070), 100U/mL Penicillin-Streptomycin (Thermo Fisher Scientific, 15140122) plus 1ug/ml α-amanitin (Sigma A2232) and 1ug/ml Puromycin (Sigma P8833). CMV clone 5 cells were nucleofected with 0.2μg tdPCP-Halo and 0.5μg TetR-RFP plasmids (Amaxa kit VCA-1003, Lonza) and seeded with media containing α-amanitin and Puromycin. 1-3 days post-nucleofection cells were stained with JF646-SNAP-tag ligand (JF646-BG) and used for imaging experiments. For staining, cells were incubated with media containing 1μM JF646-BG for 1hr, rinsed once with new media, and replaced with new media containing drugs.

#### Cell fixation

Cells were fixed with freshly-prepared 4% v/v methanol-free Formaldehyde (Thermo Scientific 28906) in 1×PBS at room temperature for 10 min, and then rinsed 3 times with 1×PBS. After fixation, samples were stored at 4°C.

#### Bead alignment/calibration sample

For routine alignment and calibration of the instrument, we use a mixture of 40nm spheres (Thermo Fisher Scientific, TransFluoSpheres 488/645, T10711) as well the 0.1μm Tetraspek beads deposited on gold EM grids with Formvar films. The 0.1μm beads provide higher signal and can facilitate fast mapping of the LLS excitation profile, while the 40nm spheres are used for fine-tuning the interferometer alignment.

### Data acquisition and analysis

3D-iLLS and conventional LLS imaging is performed by scanning the sample through the stationary LLS excitation pattern. We scan the sample simultaneously in *yz* along an axis ∼35° relatively to the LLS propagation direction (Supplementary Figure 3**c**). The *z* step size is 20nm. 3D particle tracking using modulation interferometry is performed by keeping the sample stationary and stepping the piezoelectric phase shifter in increments corresponding to relative phase changes of 90° or 60°, for 4-step and 6-step modulation cycles respectively. For far-red fluorescence emission, the step sizes are 182.5nm and 121.67nm respectively, as one modulation period corresponds to ≈730nm translation of the phase shifter. Raw image data are saved in binary format and imported for processing in MATLAB (Mathworks, 2014b).

### 3D-iLLS volume reconstruction

Raw 3D-iLLS and conventional LLS data were deskewed and deconvolved in MATLAB using the measured PSF. Deconvolution was applied to the images in Figures 2**a,b** but not to Figs. 2**c,d**. The volumetric data were imported in ImageJ for visualization and maximum projection caclulations.

### 3D-iLLS particle tracking using modulation interferometry

Raw images were imported in ImageJ and the frames from each modulation cycle were grouped together in a single maximum projection image. The maximum projection images were imported in MATLAB for 2D particle tracking analysis^20^. The trajectories of selected particles were further refined by performing a 2D Gaussian fit in 11×11 pixel ROIs to obtain more accurate *xy* coordinates. For obtaining the *z* coordinate, we first sum the intensity of the pixels in a 7×7 pixel ROI centered on the *xy* coordinate of the particle. To combine the intensities measured from Cam0 and Cam1, the intensity of Cam1 is rescaled to the same mean and standard deviation as Cam0. Finally, we calculate the phase of the intensity modulation in each cycle to extract the *z* position. In each step, the modulation phase is extracted in a 2π interval centered on the previous phase. We note that for particle displacements over successive cycles small compared to half of an interferometric period (≈265nm/2 = 132.5nm), this procedure also enables phase unwrapping and tracking over multiple interferometric fringes.

## Acknowledgements

We thank Gregory Ayzenberg (MSKCC Medical Physics) for expert machining, Daniel Mazover for assistance with CAD and Luke Lavis for dye reagents. This work was supported by a NYSTEM Postdoctoral Training Award (C32599GG; J.L.), a National Cancer Institute Grant (P30 CA008748) and by a National Institutes of Health (NIH) Director’s New Innovator Award (1DP2GM105443-01; A.P.) and by the Louis V. Gerstner, Jr. Young Investigators Fund (A.P.).

## Author Contributions

A.P. conceived, designed and supervised the study. A.P. and B.C. built the experimental apparatus. B.C. developed the data acquisition software, wrote analysis code and validated the optical performance of the 3D-iLLS setup. G.W. performed numerical calculations. J.L. prepared cell samples. A.P. performed experiments, analyzed and interpreted the data and wrote the manuscript.

## Data Availability

The data that support the findings in the paper are available from the corresponding author upon reasonable request.

## Code Availability

Custom-written code is available from the corresponding author upon reasonable request. Data acquisition and instrument control software can be requested for academic use from the corresponding author, after executing material transfer agreements with MSKCC.

