## Supplementary Material for "3D Interferometric Lattice Light-Sheet Imaging"

### Supplementary Note

**Numerical Calculations of 3D-iLLS PSFs.** We developed a numerical simulation pipeline in MATLAB that consists of the following steps: (1) estimating the excitation PSF by calculating the excitation electric field; (2) calculating the detection PSF by the electric field of single dipole emitters; (3) combining the excitation and dipole electric fields into a combined PSF, and (4) performing a near uniform orientation sampling and averaging the combined excitation-dipole electric fields of all the sampled orientations to obtain the final combined PSF. The numerical simulation pipeline illustrated with intermediate and final results is shown in Supplementary Figure 1**a**.

To calculate the desired SLM pattern for modulating the phase of the incident light and creating a certain bound 2D optical lattice we follow the procedures outlined in the original LLS publication[1]. This approach enables simulating the effects of various annular masks as well as lattice dithering. A 2D optical lattice is created by interference of light beams that exit the excitation objective lens with propagation wave vectors strictly along a cone. The excitation light thus enters the excitation objective back focal plane through an annulus (corresponding to a certain numerical aperture) of infinitesimal width. To confine the excitation light to a thin “sheet” along the x-axis by bounding the ideal 2D lattice along the z-axis, the propagation lattice wave vectors are extended along the z-axis. Therefore, the numerical simulation determines the wave vectors for a particular lattice, extends these wave vectors along *z* to confine the lattice (via the selected bounding function and the calculated SLM profile), and further constraints the *z*-extend of the wave vectors with the annular mask to achieve near non-diffracting illumination.

**Calculation of optical lattice excitation electric fields.** We start with a desired 2D optical lattice selected from the set of all five 2D Bravais lattices. The mathematical framework for calculating 2D optical lattices has been previously described[2, 3]. Below we briefly discuss the key steps involved in the numerical simulation.

First, to obtain the wave vectors for a particular 2D optical lattice, a primitive vector set **A**=[$\mathbf{a}_1$, $\mathbf{a}_2$] and its corresponding reciprocal vector set **B**=[$\mathbf{b}_1$, $\mathbf{b}_2$] (where **A** and **B** are 2 × 2 matrices) can be obtained. For each optical lattice, there are infinite sets of primitive vectors that can define it. The corresponding optical lattices are of the same type except that they exhibit different periodicities depending on the choice of **A**: the fundamental lattice of a certain type has the minimum period, while higher order sparse lattices have increasingly higher periods.

$$\mathbf{A} = [\boldsymbol{a}_1, \boldsymbol{a}_2] \tag{1}$$

For each set of primitive vectors, a corresponding reciprocal vector set **B** can be obtained.

$$\mathbf{B} = [\boldsymbol{b}_1, \boldsymbol{b}_2] = 2\pi(\mathbf{A}^T)^{-1} \tag{2}$$

A connection between the set B and the (optical) wave vectors $\{\mathbf{k}\}$ has been established by previously reported observations[2, 3]: first, a minimum of three wave vectors $\mathbf{k}_0$, $\mathbf{k}_1$, and $\mathbf{k}_2$ are required to construct a 2D optical lattice; second, these wave vectors can be constructed by

$$\mathbf{b}_n = \mathbf{k}_0 - \mathbf{k}_n, n = 1, 2 \tag{3}$$

Since the excitation beams are monochromatic, all three wave vectors are of equal length.

$$|\mathbf{k}_0| = |\mathbf{k}_1| = |\mathbf{k}_2| = \frac{2\pi}{\lambda} \tag{4}$$

By combining equations (5) and (6), a third condition can be obtained

$$\mathbf{B}^T \cdot \mathbf{k}_0 = [\boldsymbol{b}_1^T \cdot \boldsymbol{b}_1, \boldsymbol{b}_2^T \cdot \boldsymbol{b}_2]^T \equiv \frac{\boldsymbol{\beta}}{2} \tag{7}$$

With all three equations, the first wave vector $\mathbf{k}_0$ can be solved as

$$\mathbf{k}_0 = (\mathbf{B}^T)^{-1} \cdot \mathbf{B}^T \cdot \mathbf{k}_0 = (\mathbf{B}^T)^{-1} \cdot \frac{\boldsymbol{\beta}}{2} = \frac{1}{2\pi} \cdot \mathbf{A} \cdot \frac{\boldsymbol{\beta}}{2} = \frac{\mathbf{A} \cdot \boldsymbol{\beta}}{4\pi} \tag{8}$$

And the rest of the wave vectors, $\mathbf{k}_1$ and $\mathbf{k}_2$, can be solved as

$$\mathbf{k}_n = \mathbf{k}_0 - \mathbf{b}_n = \frac{\mathbf{A} \cdot \boldsymbol{\beta}}{4\pi} - \mathbf{b}_n, n = 1, 2 \tag{9}$$

By Fourier transforming these wave vectors, the initial desired 2D lattice can be derived:

$$E_{ideal;\, fundamental/sparse} = FT[(\mathbf{k}_0, \mathbf{k}_1, \mathbf{k}_2)] \tag{10}$$

However, these fundamental and sparse 2D optical lattices have broad foci that extend throughout the unit cell, thus limiting their use for creating thin sheets of illumination. To overcome this difficulty, composite 2D optical lattices are explored, which consist of a greater number of wave vectors than the minimum three: composite optical lattices have more confined excitation foci due to the constructive interference of the additional wave vectors. One way of generating more wave vectors is to perform symmetry operations on the initial three wave vectors. The maximum number of wave vectors is obtained through the maximum number of allowed symmetry operations, generating a maximally symmetric (composite) 2D optical lattice.

$$E_{ideal;composite\ (max\ symm)} = FT[symmetry\_operation(\mathbf{k}_0, \mathbf{k}_1, \mathbf{k}_2)] \tag{11}$$

We can confine the ideal 2D lattice along the z-axis into a lattice light sheet using an arbitrary bounding function $\psi(z)$.

$$E_{bound} = \psi(z) \cdot Re(E_{ideal}) \tag{12}$$

The profile of this bound lattice is then used to create the phase pattern of the binary SLM with a Heaviside step function H.

$$\varphi_{SLM} = \pi \cdot H(E_{bound} - \varepsilon) \tag{13}$$

where $\varepsilon$ is an arbitrary cutoff.

Once the phase pattern for the SLM is obtained, we derive the excitation profile at the *xz* focal plane using an annular mask *N* that removes unwanted diffractions after transforming the phase-modulated beam.

$$PSF_{ex} = |FT\left[N \cdot FT\left(e^{i\varphi_{SLM}}\right)\right]|^2 \tag{14}$$

Although the above 2D simulation reveals the excitation profile at the *xz* focal plane, it does not describe how the bound lattice propagates along the y-axis, which ultimately determines the effective field of view. We calculate the 3D excitation electric field, which is approximately expressed in the near-focus space as[4]:

$$\vec{E}(x,y,z) = (E_x, E_y, E_z) \tag{15}$$

$$\begin{cases} E_x = -iA(I_0 + I_2 cos2\varphi) \\ E_y = -iAI_2 sin2\varphi \\ E_z = -2AI_1 cos\varphi \end{cases}$$

where $I_0$, $I_1$, and $I_2$ are integrals over the aperture of the excitation objective; $A$ is a scalar; $\varphi$ is the azimuth in the cylindrical coordinate system.

However, numerically solving the above equations is not efficient due to the inherent nested loops used to calculate the integrals. Interestingly, an alternative implementation of the integrals as a Fourier transform significantly increases the speed of numerical calculations[5] and thus is incorporated in this simulation.

**Caclulation of single dipole emission electric fields.** After calculating the electric field of the excitation LLS in 3D, we next simulate the 3D electric field of the emission from a single dipole. The dipole emission imaged by one of the detection objectives can be expressed as:

$$\mathbf{E}_1(r, z, \varphi) = (E_x, E_y, E_z) \tag{16}$$

$$E_a = B \int_0^{\theta_{max}} d\theta sin\theta \cdot G_{ab}^E(r, \theta, \varphi) p_b e^{ikzcos\theta} \tag{17}$$

where $r$ is the distance from the optical axis ($z$-axis for emission detection) to the point-of-interest; $\theta$ is the angle between the vector pointing from the focus to the point on the aperture; $\theta_{\max}$ is the maximum angle of the aperture of the detection lens; $\varphi$ is the azimuth angle in cylindrical coordinates: z is the distance away from the focus along the optical axis; $G_{ab}^E$ is a tensor whose components are given by Ref 6.

When the apertures of two opposite imaging objectives are superimposed coherently, the dipole electric field can be combined as:

$$\mathbf{E}_{dipole} = \mathbf{E}_1 + \mathbf{E}_2 \tag{18}$$

$$\mathbf{E}_2(r, z, \varphi) = \begin{pmatrix} 1 & 0 & 0 \\ 0 & -1 & 0 \\ 0 & 0 & -1 \end{pmatrix} \mathbf{E}_1(r, -z, -\varphi) \mathrm{e}^{\Delta\Psi} \tag{19}$$

where $\Delta\Psi$ denotes the path-length difference between the two detection interferometer arms.

**Calculation of the overall PSF.** Once the electric fields of the LLS excitation and the dipole emission are obtained, the response of the interferometric LLS microscope to a single point emitter of a particular dipole orientation can be described with the overall point spread function (PSF). In dithered mode, the *x*-axis continuous illumination is achieved by using the *x*-galvo to scan the excitation beam over multiples of that lattice period along the *x*-axis, which can be modeled numerically by shifting the 3D electric field over one period. Therefore, the averaged overall PSF is calculated as:

$$PSF_{overall,i} = |\mathbf{E}_{dipole,i}|^2 \cdot \int_x^T dx \cdot |\mathbf{E}_{ex} \cdot \mathbf{p}_{dipole,i}|^2 \quad (20)$$

where $\mathbf{p}_{\text{dipole,i}}$ is the i-th orientation of a particular emitting dipole; T is the period of the 2D optical lattice along the *x*-axis.

We further perform a near uniform sampling of points across a sphere to account for possible orientations explored by organic dyes or fusion fluorescent proteins that are either conjugated with a flexible linker or simply freely diffusing in the cell. However, for N other than 2, 3, 4, 6, 8, 10, or 12, there is no analytical solution to place N points at equal distance to the adjacent points on a spherical surface. Random uniform sampling of z in [-1, 1] and φ in [0, 2π] in a spherical coordinate system introduces clustering[7], which is more pronounced when N is relatively small. To avoid potential bias in the numerical calculations of the overall PSF, we follow an alternative sampling method that results in near uniform distribution of dipole orientations[8]: the sphere is first separated into equally-spaced equators, each of which is then separated into segments of length approximately equally to the inter-equator distance with ends being the sampled orientations (Supplementary Figure 1**b**).

Finally, we sum the overall PSFs of all sampled orientations to generate the averaged overall PSF:

$$PSF_{overall} = \sum_{i}^{n} PSF_{overall,i} \tag{21}$$

The results for calculated 3D-iLLS PSFs for different types of bound 2D lattice excitation profiles are shown in Supplementary Figure 2.

**Supplementary Figures & Legends**

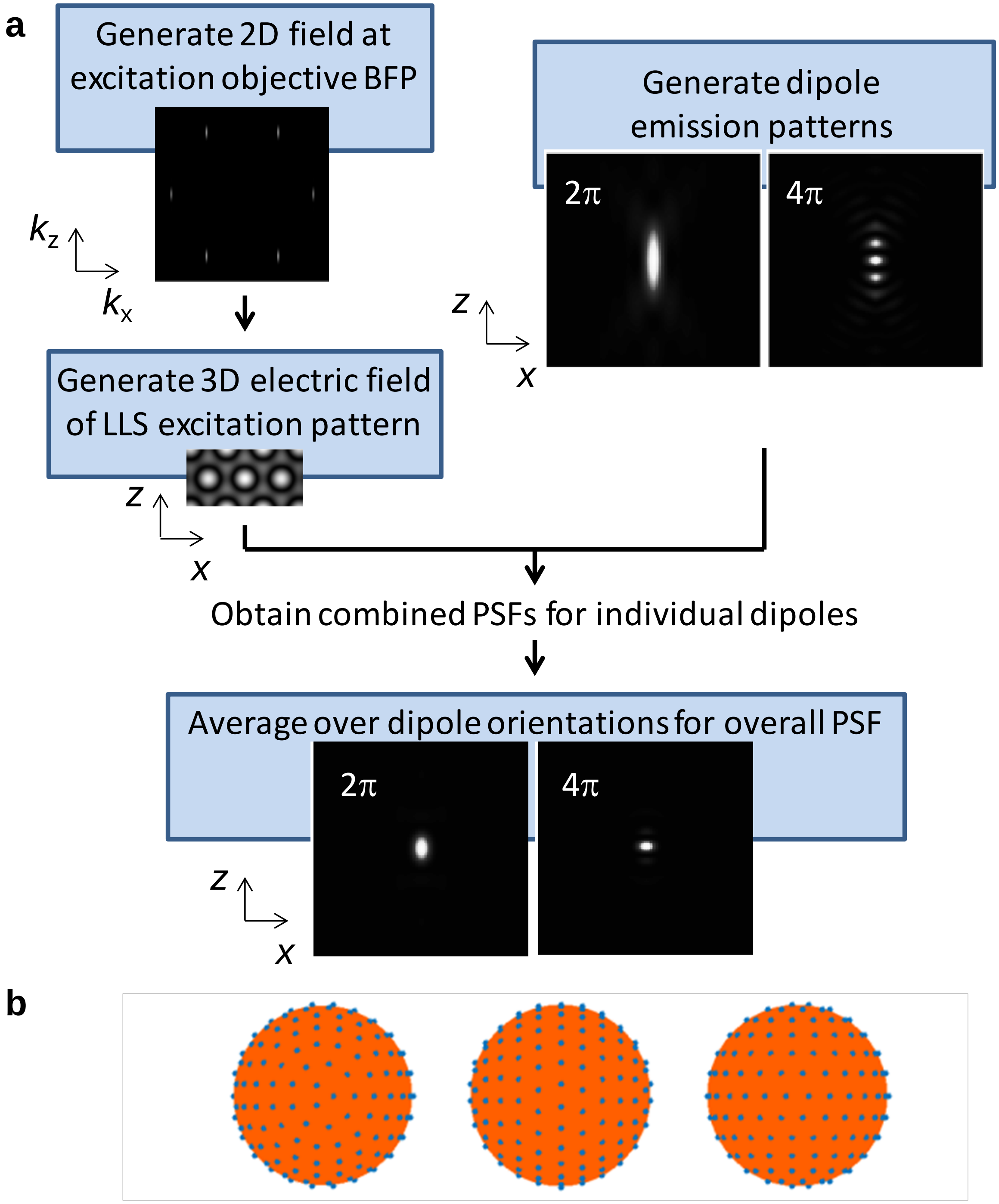


Supplementary Figure 1 **Numerical calculation of 3D-iLLS PSFs. a,** Simulation pipeline. **b,** Near uniform sampling of 214 orientations viewed from three angles.

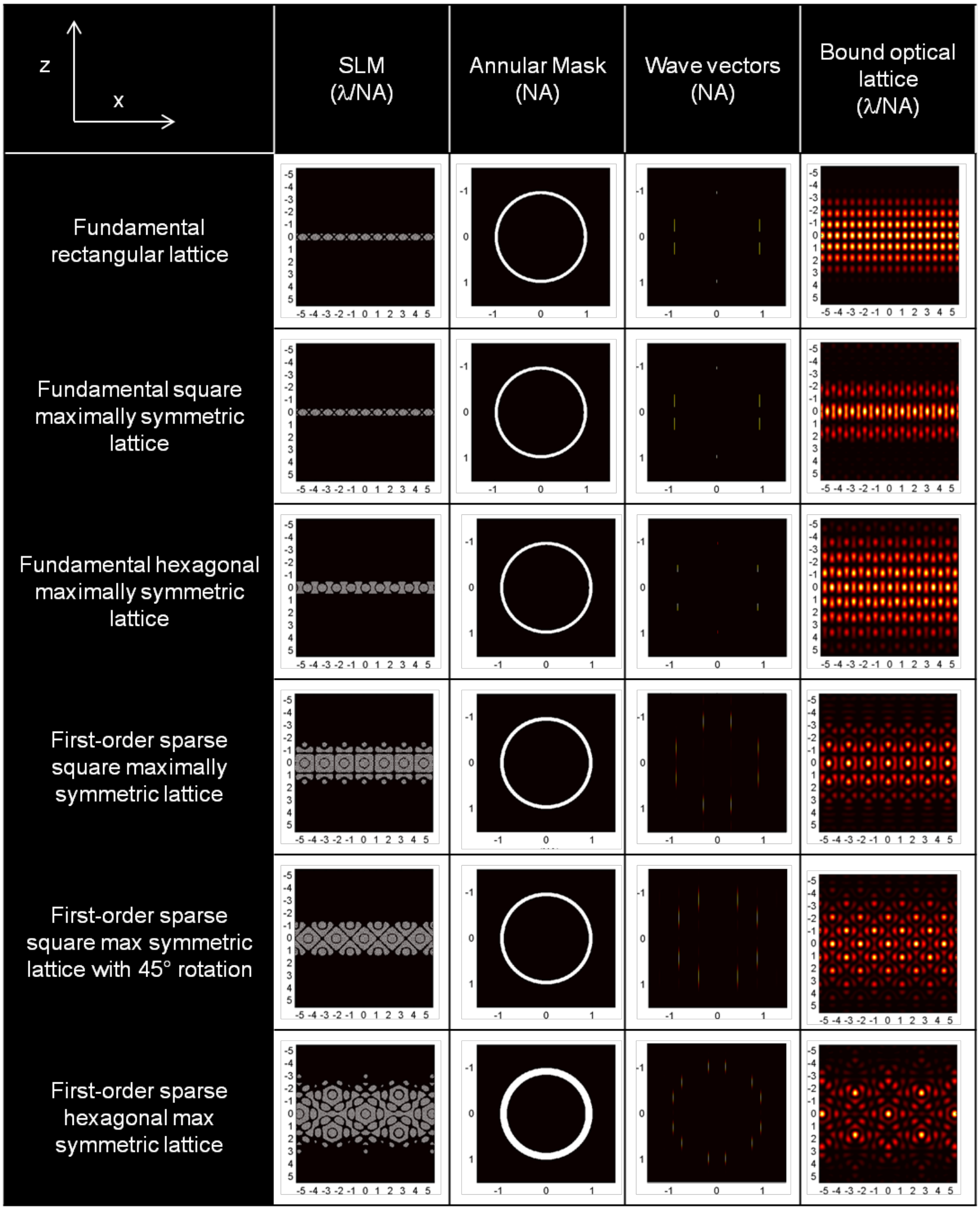
z
x
SLM
(λ/NA)
Annular Mask
(NA)
Wave vectors
(NA)
Bound optical
lattice
(λ/NA)
Fundamental rectangular lattice
Fundamental square maximally symmetric lattice
Fundamental hexagonal maximally symmetric lattice
First-order sparse square maximally symmetric lattice
First-order sparse square max symmetric lattice with 45° rotation
First-order sparse hexagonal max symmetric lattice

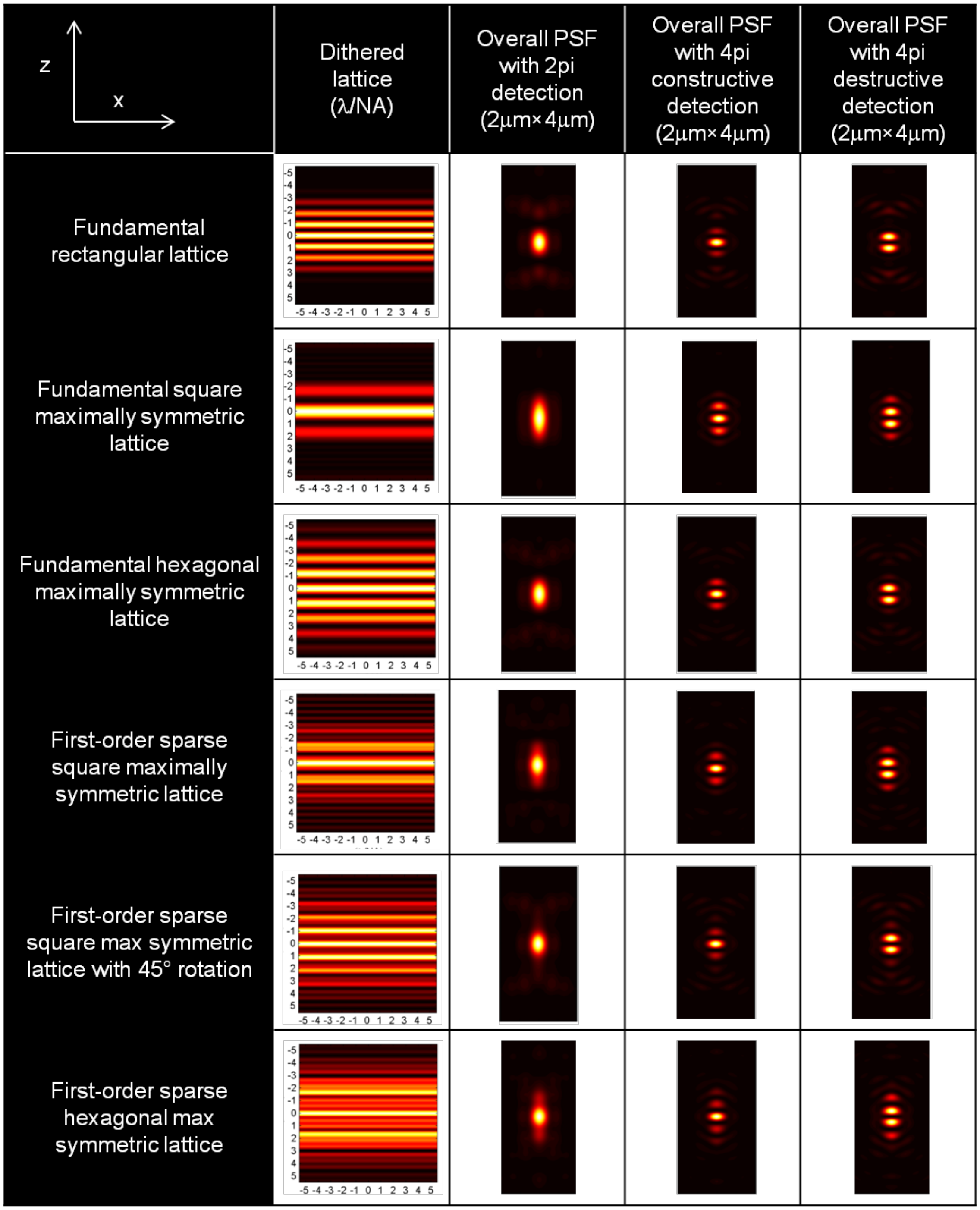


Supplementary Figure 2 **Numerical 3D-iLLS PSFs for different excitation lattices and comparison with conventional LLS.** Simulation parameters are given in Supplementary Tables 1 and 2.

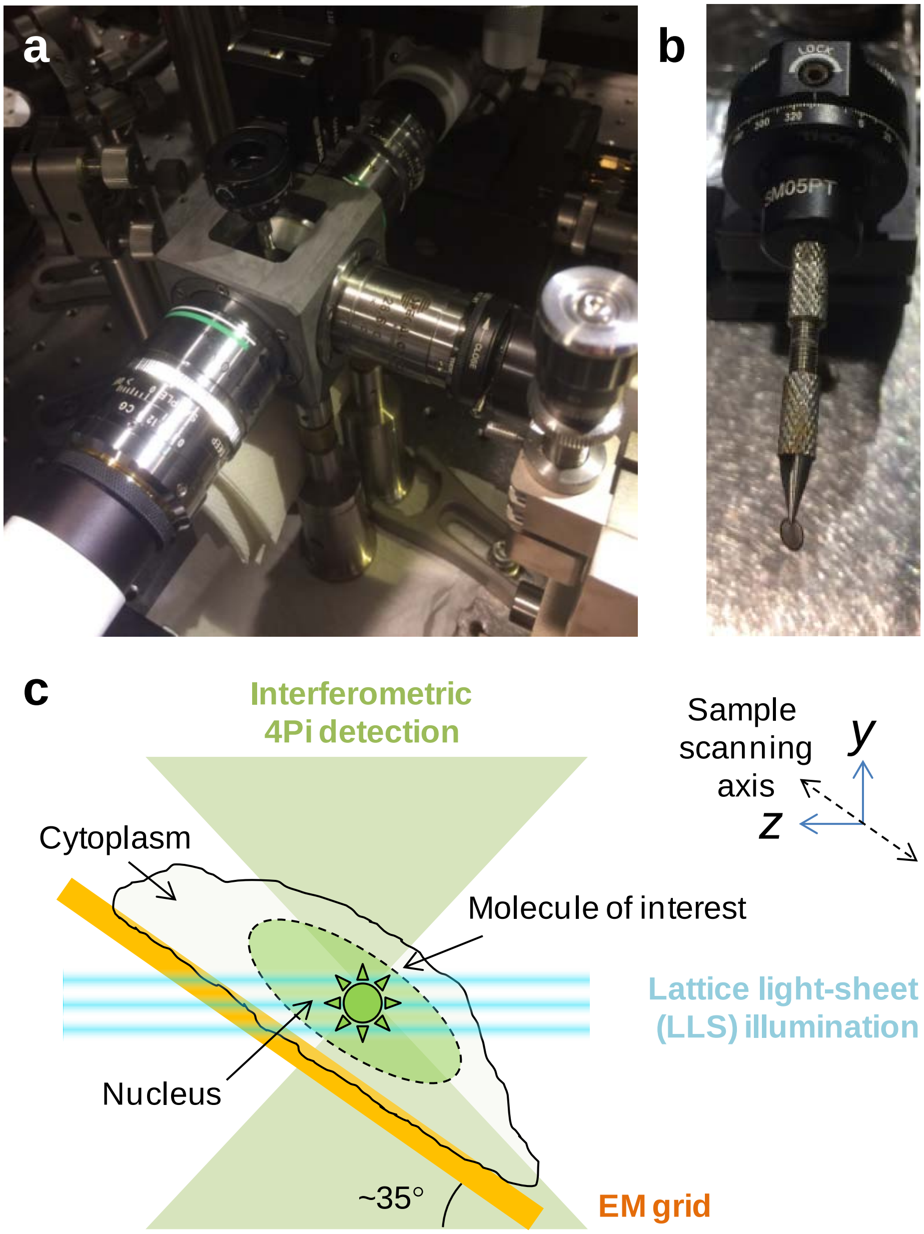


Supplementary Figure 3 **Three-objective 3D-iLLS configuration, liquid sample cell, sample holder and sample mounting geometry. a,** photograph of 3D-iLLS setup, highlighting the three-objective configuration, the liquid sample cell and the top-immersion sample holder. **b,** photograph of the pincher-grip sample holder with mounted EM grid. **c,** sample mounting geometry.

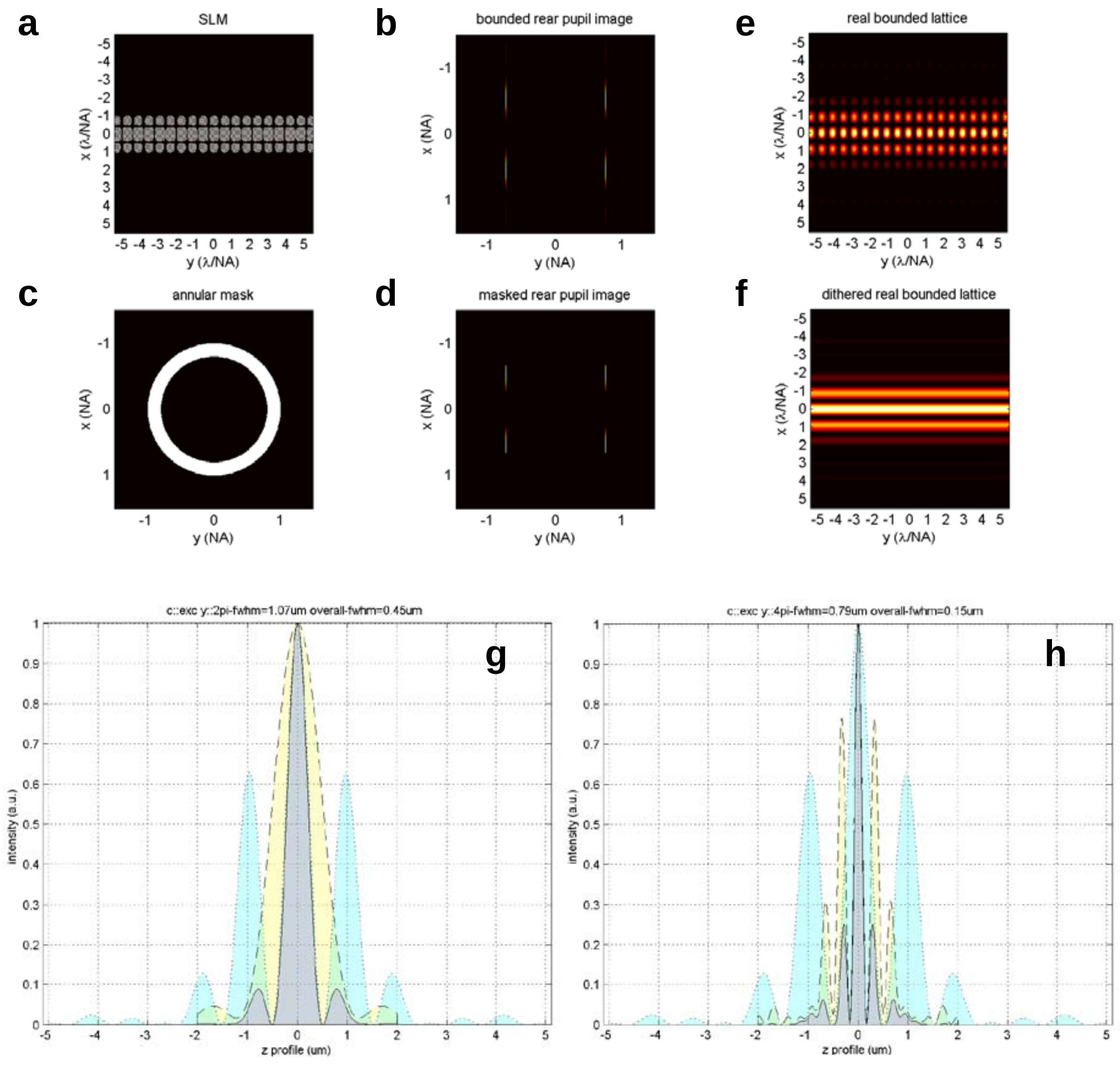


Supplementary Figure 4 **Optimized 3D-iLLS PSF based on a fundamental rectangular 2D bound lattice. a,** SLM pattern and **b,** corresponding intensity at rear pupil. **c,** Annular mask. **d,** Intensity at rear pupil after annular mask. **e,** Resulting 2D bound lattice in real space and **f,** corresponding dithered lattice excitation pattern. **g,** axial profile of conventional LLS PSFs. **h,** Axial profile of 3D-iLLS PSFs. Cyan: excitation; yellow: detection; gray: overall.

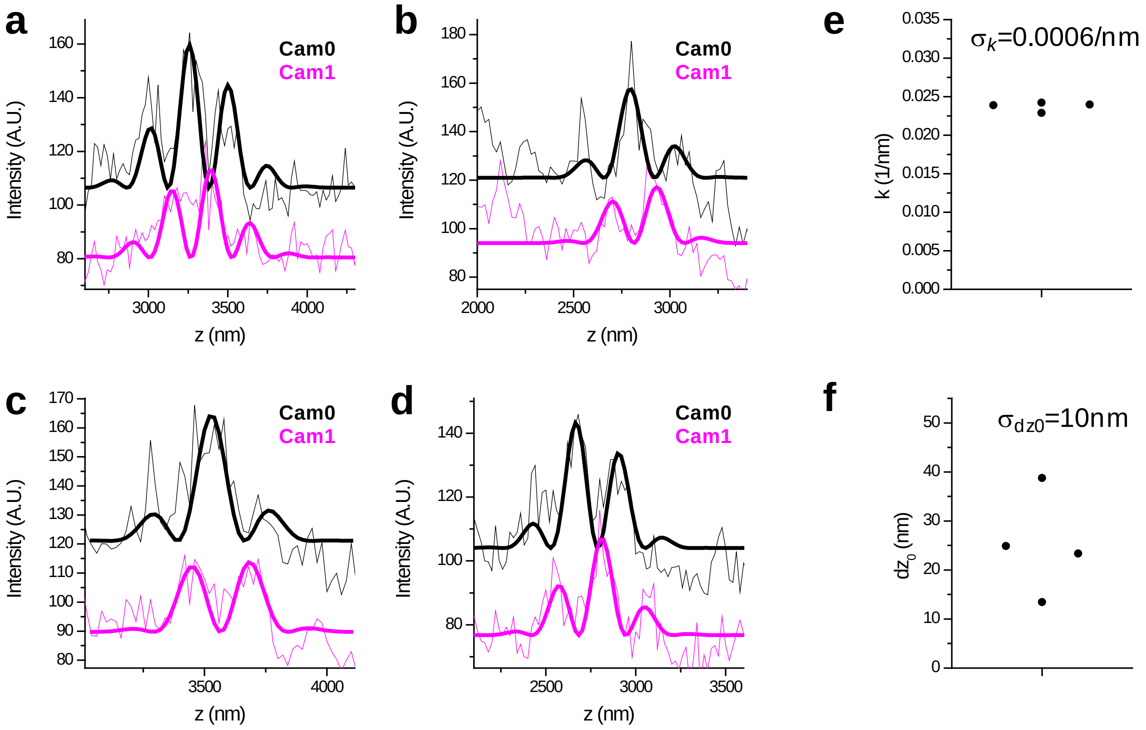


Supplementary Figure 5 **Localization of Brd4 clusters in reconstructed 3D-iLLS images. a-d,** axial profiles of individual Brd4 clusters. Thick lines show non-linear least-squares fits to equations of the form $B+\frac{A}{2}(1\pm\cos(k(z-z_0)+\theta))e^{-\frac{(z-z_0)^2}{2\sigma_z{}^2}}$ for Cam0 and Cam1, respectively. Global fitting is performed, with shared $k, \theta$ and $\sigma_z$ parameters. We obtain two separate localization measurements of the parameter $z_0$ that indicates the center position of the cluster, estimated independently from Cam0 and Cam1. **e,** oscillation wave-vector $k$ is 0.02375±0.00059 $nm^{-1}$ (mean±SD), indicating a relative error $\sigma_k/k$ of ≈2.5%. **f,** the center position $z_0$ shows a systematic offset between the two cameras of $dz_0$=25nm and an r.m.s localization error $\sigma_{dz0}$=10nm. These systematic and random errors relative to the oscillation period ($2\pi/k$=265nm) are ≈9% and ≈4% respectively.

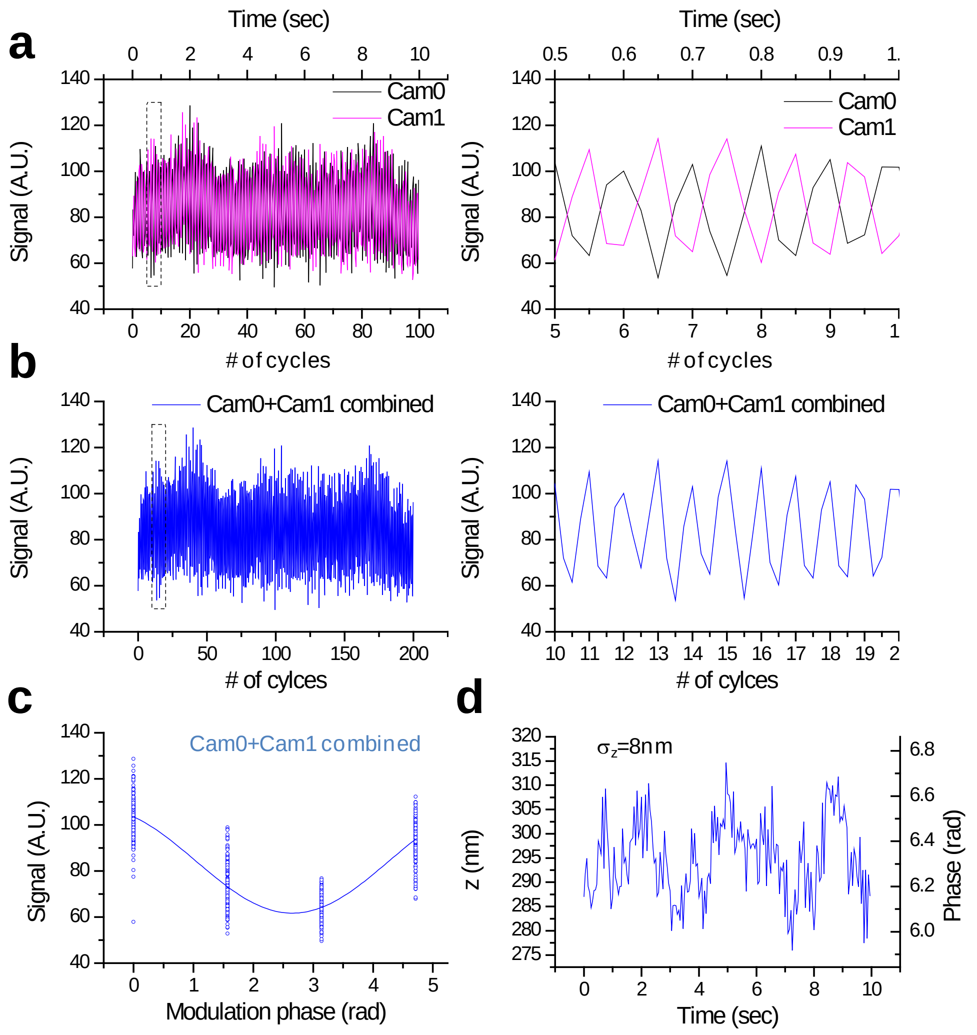


Supplementary Figure 6 **Axial localization performance with 3D-iLLS and 4-phase modulation interferometry. a,** Signals from a 40nm bead on Cameras 0 and 1, over 100 4-step modulation cycles. The piezoelectric phase shifter is stepped in 182.5nm increments, corresponding to 0°, 90°, 180° and 270° relative phases. Right panel: zoom-in of the dotted region in the left panel, illustrating the anti-correlated signal modulation of Cam0 vs. Cam1. **b,** Signals from Cam0 and Cam1 are combined into a single modulation cycle, doubling the temporal resolution. Right panel: zoom-in of the dotted region in the left panel. **c,** Superposition of all 200 modulation cycles by collapsing the *x* axis in the interval

[0-2π), showing excellent stability and reproducibility of the setup. Solid line: fit to a sine wave. **d,** Extracted phase and *z* coordinate, showing $\sigma_z \approx 8$nm r.m.s. localization precision.

**Supplementary Table 1. Lattice-type-specific parameters used in simulating the six different 2D lattice light sheet excitation profiles in Supplementary Figure 1.**

| 2D lattice | Bounding Gaussian function σ (λ/NA) | Wave vector length (NA) | Annular mask inner diameter (NA) |
|---|---|---|---|
| Fundamental rectangular | 1.5 | 90% | 80% |
| Fundamental square maximally symmetric | 0.15 | 92% | 93% |
| Fundamental hexagonal maximally symmetric | 0.3 | 100% | 93% |
| First-order sparse square maximally symmetric | 0.8 | 94% | 92% |
| First-order sparse square maximally symmetric with 45° rotation | 0.8 | 94% | 92% |
| First-order sparse hexagonal maximally symmetric | 1.5 | 100% | 85% |

**Supplementary Table 2. Common parameters used in simulating the six different 2D lattice light sheets in Supplementary Figure 1.**

| Common simulation parameters | Values |
|---|---|
| Excitation wave length | 641 nm |
| Heaviside coefficient ε | 0.1 |
| Annular mask outer diameter | 100% NA |
| Excitation objective NA | 0.65 |
| Excitation objective focal length | 7 mm |
| Refractive index of imaging medium | 1.33 |
| Emission objective wave length | 700 nm |
| Emission objective NA | 1.1 |
